# Optimization of avian perching manoeuvres

**DOI:** 10.1101/2021.08.28.458019

**Authors:** Marco KleinHeerenbrink, Lydia A. France, Caroline H. Brighton, Graham K. Taylor

**Affiliations:** Department of Zoology, University of Oxford, OX1 3SZ, UK; The Alan Turing Institute, London, NW1 2DB, UK

## Abstract

Perching at speed is amongst the most challenging flight behaviours that birds perform^1,2^, and beyond the capability of current autonomous vehicles. Smaller birds may touchdown by hovering^3-8^, but larger birds typically swoop upward to perch^1,2^ – presumably because the adverse scaling of their power margin prohibits slow flapping flight^9^, and because swooping transfers excess kinetic to potential energy^1,2,10^. Perching is risky in larger birds^6,11^, demanding precise control of velocity and pose^12-15^, but it is unknown how they optimize this challenging manoeuvre. More generally, whereas cruising flight behaviours such as migration and commuting are adapted to minimize cost-of-transport or time-of-flight^16^, the optimization of unsteady flight manoeuvres remains largely unexplored^7,17^. Here we show that swooping minimizes neither the time nor energy required to perch safely in Harris’ hawks *Parabuteo unicinctus*, but instead minimizes the distance flown under hazardous post-stall conditions. By combining motion capture data from 1,563 flights with flight dynamics modelling, we found that the birds’ choice of where to transition from powered dive to unpowered climb minimizes the distance from the landing perch over which very high lift coefficients are required. Time and energy are therefore invested to maintain the control authority needed to execute a safe landing, rather than being minimized continuously as in technical applications of autonomous perching under nonlinear feedback control^13^ and deep reinforcement learning^18,19^. Naïve birds learn this behaviour on-the-fly, so our findings suggest an alternative reward function for reinforcement learning of autonomous perching in air vehicles.

---

The exquisite perching performance of birds has inspired many efforts to achieve perching capability in autonomous aircraft^10,13,14,18,20-24^, which is made especially challenging by the lack of a runway to bleed speed after landing. This creates a precise targeting requirement that is made even more demanding by the difficulty of maintaining control authority at the low airspeeds needed before touchdown^10,13,20^. Although some kinetic energy is converted to gravitational potential energy when climbing up to a perch^1,2,10^, most is either lost through aerodynamic drag or dissipated upon impact^3,7,25-27^. Aerodynamic braking is therefore crucial to avoiding a dangerously energetic collision, but the high angles of attack that this requires compromise flight control as the wing stalls^10,13,20,24^. Birds execute a characteristic rapid pitch-up manoeuvre when perching, which transiently increases aerodynamic force production^28^ and is expected to delay stall^1,2,6,22,24,29^. The rapidity of this manoeuvre makes planning of its entry conditions critical^18^, which begs the question of how the flight trajectory and control inputs are optimized. More generally, these perching dynamics provide a tractable test case for asking how animals optimize complex unsteady motions.

To address these questions, we rigged a large custom-built motion capture studio to record *n* = 4 captive-bred Harris’ hawks flying between perches for food (see Methods; Supplementary Movie 1). The hawks wore retroreflective markers enabling us to reconstruct their flight trajectories at a 120 or 200 Hz sampling rate (Fig. 1). Three of the birds were juvenile males that had only flown short distances previously; the other was an experienced adult female. We collected trajectory data from 1,585 flights at 5, 7, 9, or 12 m perch spacing and 1.25 m perch height, after an initial familiarisation period comprising 100 flights per bird made at 12 m perch spacing. Perch spacing was held at 12 m for the 2-3 weeks following, to allow us to confirm the stability of the behaviour, and was subsequently randomized daily at 5, 7, or 9 m. The naïve birds used flapping to fly directly between the perches for the first few flights of their familiarisation period (Fig. 2A), but soon adopted the indirect swooping behaviour characteristic of experienced birds (Fig. 2B-E). Swooping was initiated by jumping forwards into a dive involving several powerful wingbeats, which transitioned into an unpowered climb comprising a rapid pitch-up manoeuvre that ended with the body almost vertical and with the wings outstretched as the feet contacted the perch (Fig. 1). Climbing was mainly executed by gliding, with occasional half-flaps constituting the ventral part of a flapping motion, which we interpret as corrective control inputs rather than wingbeats supplying thrust to offset drag.

**Figure 1.**
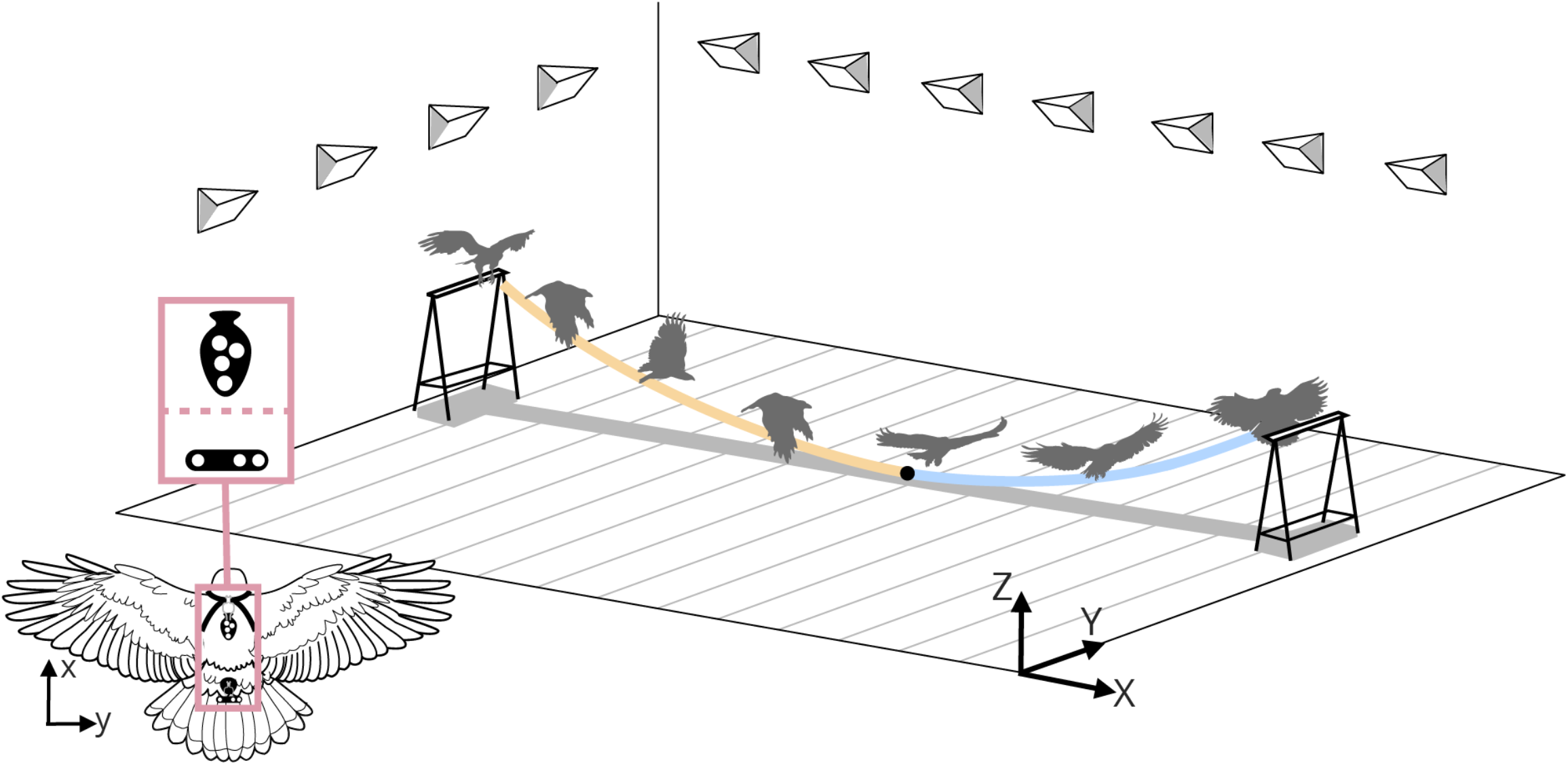
Schematic of characteristic swooping trajectory and data acquisition. Harris’ hawks were flown between perches in a purpose-built motion capture studio, wearing a template of retroreflective markers close to the centre of mass (inset panel; tail markers also shown). Swooping was initiated by a take-off jump, followed by a powered dive (yellow line). This transitioned at its lowest point (black dot) into an unpowered climb (blue line), involving a rapid pitch-up manoeuvre ending with the body almost vertical, and the wings outstretched as the feet contacted the perch.

**Figure 2.**
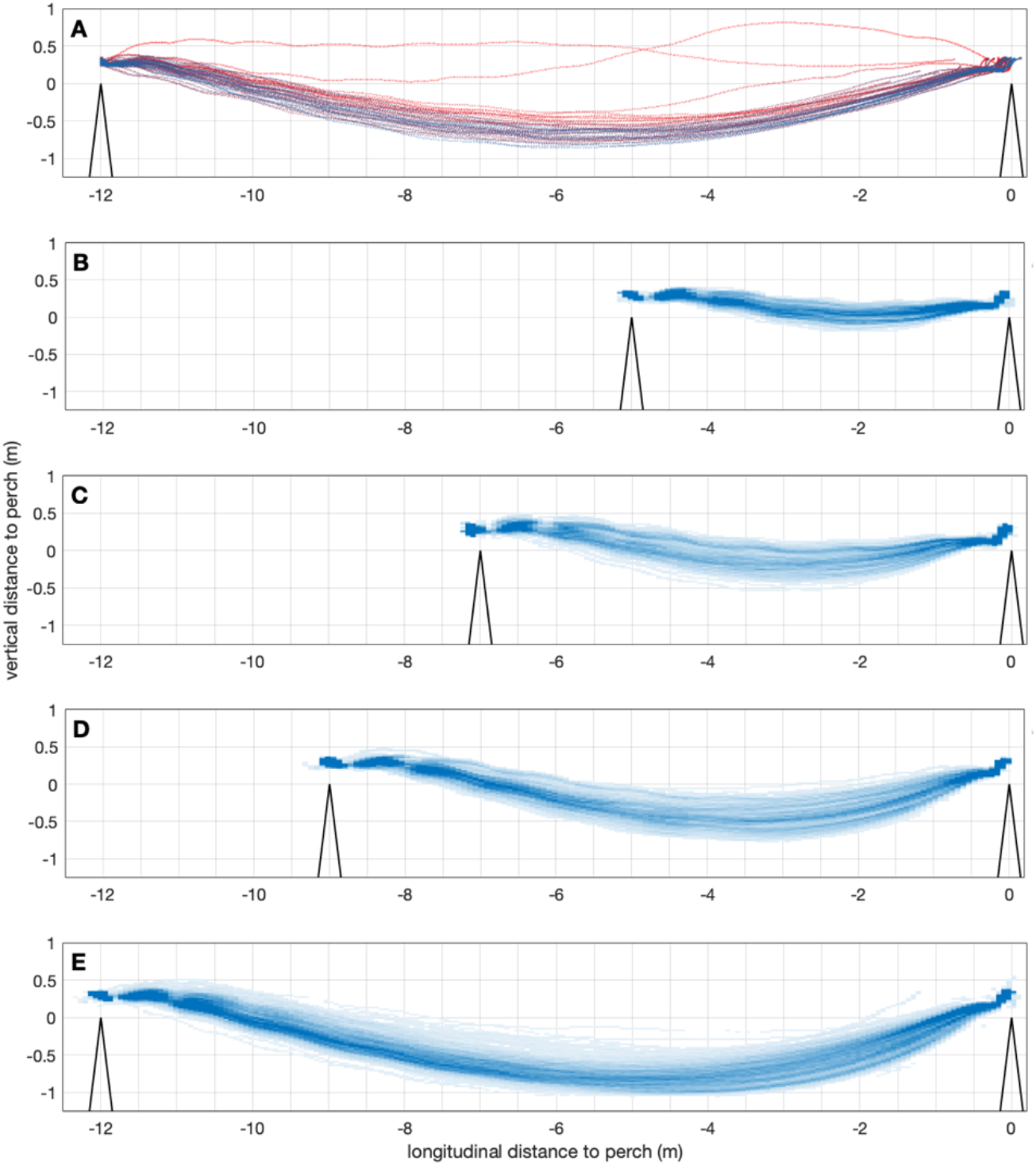
Measured swooping trajectories of perching Harris’ Hawks. **(A)** Ontogeny of 45 flight trajectories recorded at 12 m perch spacing during the initial familiarisation period for juvenile bird “Toothless”; earlier flights in red; later flights in blue. Note the more direct trajectory taken on earlier flights, and quick acquisition of a swooping trajectory characteristic of experienced birds. (**B-E**) Spatial histograms showing pooled trajectory data from 8 = 1,563 flights for all *n* = 4 hawks at (B) 5 m, (C) 7 m, (D) 9 m, and (E) 12 m perch spacing. These spatial histograms do not include trajectories recorded during the initial familiarisation period.

We summarized the geometry of each observed trajectory by measuring the location of its point of minimum height as a proxy for the location of the transition from powered dive to unpowered climb. We used a linear mixed effects model to characterize how this geometry varied with perch spacing across individuals, excluding 22 outliers with high residual error (see Methods). The relative longitudinal position of the observed transition point (marginal mean ± s.e. at mean perch spacing: 61.0 ± 1.20 % of perch spacing distance) did not vary significantly in relation to perch spacing over the range of distances tested (slope estimate ± s.e.: -0.078 ± 0.15 % m^−1^; *t*_(1561)_=-0.52; *p*=0.60). This finding was robust to the re-inclusion of the 22 outliers that we had excluded from this and subsequent analyses. In contrast, the relative depth to which the birds dived (marginal mean ± s.e. at mean perch spacing: 3.20 ± 0.49 % of perch spacing distance) increased significantly with perch spacing (slope estimate ± s.e.: 1.06 ± 0.071 % m^−1^; *t*_(1561)_=14.9; *p*<0.0001). In each case, the consistency with which different individuals adopted qualitatively similar swooping behaviour at different perch spacing distances (Fig. 2) suggests that they may have acquired this through individual learning optimizing some common performance objective. What might this objective function be?

Flying between perches is energetically demanding because of the high aerodynamic power requirements of slow flight, and our hawks were usually panting visibly by the end of a trial. Guided by previous work on perching parrotlets^7^, we hypothesized that the hawks would have learned trajectories minimizing the energetic cost of flying between the perches. An alternative hypothesis is that they learned trajectories minimizing the time of flight^16^, which would make sense for a predator adapted to exploit fleeting feeding opportunities^17^, and would maximize the net rate of energy gain when flying at speeds below the minimum power speed^16^. Could optimization of either objective explain the swooping behaviour that we observed? Diving exploits gravity to reach higher speeds quicker^30^, so a swooping flight path might be expected to reduce flight duration, analogous to the brachistochrone problem in which a curved path minimizes the time of travel for a particle falling under gravity between two points spaced vertically and horizontally^31^. Diving might also be expected to reduce the energy required for flight, by raising the bird’s airspeed closer to its minimum power speed^30^. It therefore seems intuitive that swooping could reduce both the energetic cost and time of flight.

We used a simplified flight dynamics model to quantify how these two performance objectives were influenced by the birds’ behaviour, comparing the trajectories predicted to minimize time and energy with the birds’ observed trajectories. We modelled perching as a two-phase flight behaviour, comprising a powered dive switching to an unpowered glide. Our simulations incorporated inter-individual variation in body mass, wingspan, wing area, and take-off speed (Table 1), and as a first-order modelling approach, we assumed that aerodynamic lift and power were held constant for each phase. We determined thrust as the ratio of aerodynamic power to airspeed, and modelled aerodynamic drag by a theoretical drag polar parameterized using wind tunnel measurements from Harris’ hawks^32^. Ground effect is expected to enhance aerodynamic efficiency when flying close to a surface^33,34^, but the birds only dived close to the ground at 12 m perch spacing (Fig. 2E), and for so brief a period that this had little effect on the predicted flight trajectory (Ext. Fig. 1) when modelled aerodynamically^34^. We therefore ignore ground effect in the optimization analysis, which greatly simplifies its implementation and interpretation. The resulting model captures the indirect swooping flight behaviour of experienced birds and – at shallow dive angles – the direct flight behaviour of naïve birds.

**Table 1.** Measurements and model parameters by bird.

| | sex | age | $m$ (kg) | $b$ (m) | $S$ (m<sup>2</sup>) | $\bar{V}_0$ (m s<sup>-1</sup>) | $\bar{V}_{\text{end}}$ (m s<sup>-1</sup>) | $\hat{P}_{\text{dive}}/m$ (W kg<sup>-1</sup>) |
| --- | --- | --- | --- | --- | --- | --- | --- | --- |
| Drogon | ♂ | juv. | 0.660 | 1.01 | 0.1895 | 3.9 | 2.3 | 23.2 |
| Rhaegal | ♂ | juv. | 0.620 | 1.02 | 0.1918 | 4.0 | 2.5 | 18.9 |
| Ruby | ♀ | ad. | 0.874 | 1.08 | 0.2146 | 3.9 | 2.3 | 22.7 |
| Toothless | ♂ | juv. | 0.738 | 1.07 | 0.2098 | 3.8 | 2.5 | 23.0 |

juv. juvenile; ad. adult; $m$ total mass; $b$ wingspan; $S$ wing area; $\bar{V}_0$ mean observed take-off speed; $\bar{V}_{\text{end}}$ mean observed landing speed; $\hat{P}_{\text{dive}}/m$ best-fitting specific power setting.

For a given constant power setting (*P*_dive_), every modelled flight trajectory is therefore parameterized by the initial dive angle (*γ*_0_) and constant lift setting (*L*_dive_) for the powered phase. The initial conditions for the unpowered phase are given by the speed (*V*_T_) and position (*x*_T_,, _T_) of the bird at the transition point where it switches from powered to unpowered flight. Given these entry conditions, the lift setting for the unpowered phase (*L*_glide_) is uniquely determined by the constraint that the trajectory must intercept the perch at the mean landing speed observed for each bird (Table 1). By enforcing this constraint, we were able to identify a set of feasible parameter settings that would bring the bird safely to its perch. Specifically, for a given constant power setting, *P*_dive_, the set of all feasible pairs {*γ*_0_, *L*_dive_} maps onto a line of feasible transition points {*x*_T_, *y*_T_}, each associated with a corresponding transition speed *V*_T_ (Fig. 3; Ext. Fig. 2). This convex curve includes transition points characterising a range of feasible perching behaviours, varying from almost level flight involving a short, powered phase and long unpowered phase (Fig. 3A), to almost level flight involving a long, powered phase and minimally short unpowered phase (Fig. 3B), with intermediate transition points being associated with swooping trajectories resembling those of experienced birds (Fig. 3C). Minimizing the sum of the squared distance between the locations of the observed transition points and the line of feasible transition points predicted under the model yielded specific power estimates ranging from 18.9 to 23.2 W kg^−1^ (Table 1), which is well below the maximum available power that Harris’ hawks have at their disposal when climbing^35,36^.

**Figure 3.**
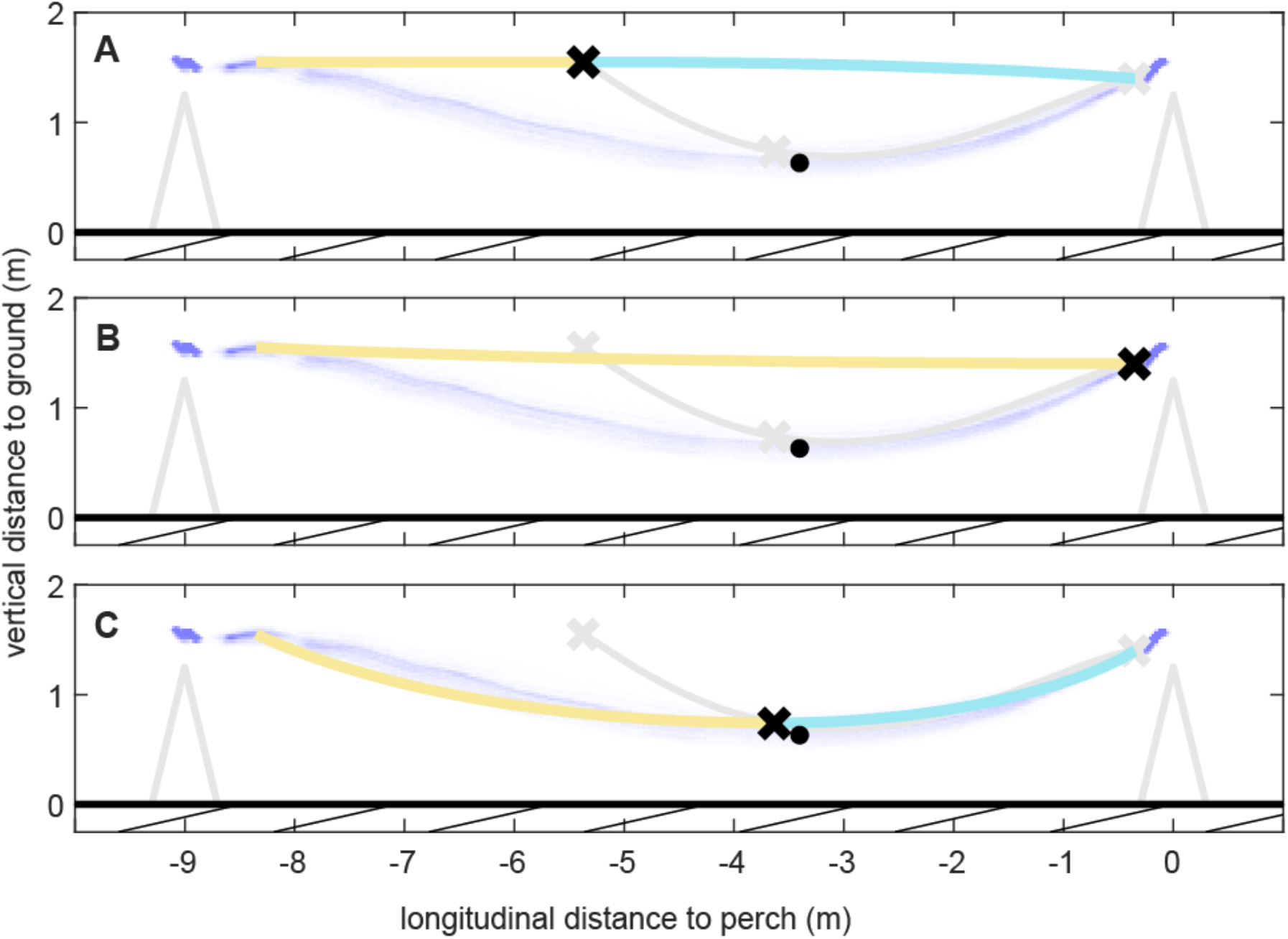
Optimal perching trajectories minimizing different cost functions. Solid lines show trajectories predicted for juvenile bird “Drogon” at 9 m perch spacing under the two-phase perching model, comprising a powered dive (yellow line), transitioning into an unpowered glide (blue line), minimizing: **(A)** energetic cost; **(B)** flight duration; or **(C)** stall distance. The corresponding optimal transition point (black cross) along the line of feasible transition points (grey line) is only predicted to fall close to the observed transition point (black dot) at the bottom of the flight trajectory if stall distance is optimized (C). Each panel shows the optimal transition points under the alternative objective functions as grey crosses; observed trajectories are plotted as a spatial histogram (blue shading).

The trajectories modelled at the best-fitting power setting and mean landing speed for each bird reveal some unexpected features. First, although diving more steeply allows faster speeds to be reached sooner on the powered phase, shortening the unpowered phase proves more effective in reducing flight duration. The time-optimal solution therefore involves a shallow powered dive and minimally short unpowered phase, with the transition point falling close to the landing perch (Fig. 3B). Second, although the efficiency of lift production is enhanced at faster airspeeds, more energy is needed in a deeper dive because of the higher lift needed to swoop up to the perch. The energy-optimal solution is therefore an almost level flight trajectory, with the transition point falling towards the take-off perch, and a long shallow glide in which airspeed is lost continuously to maintain altitude (Fig. 3A). It follows that neither time nor energy minimization straightforwardly explains the deep swooping behaviour that our birds acquired.

It is possible in principle that our birds made a particular trade-off between time and energy minimization, leading to the selection of an intermediate transition point involving swooping, but it is reasonable to ask whether they might have been optimizing a different objective function altogether. Minimizing either time or energy requires the bird to sustain very high lift coefficients, *C*_*L*_ = 2*L*/(*ρV*^2^*S*), where *L* is lift, *ρ* is air density, *V* is airspeed, and *S* is wing area. These are needed to provide high braking force when minimizing flight time (Fig. 3B), and to support body weight at ever-decreasing airspeeds when minimizing energetic cost (Fig. 3A). Very high lift coefficients can occur transiently during perching^37,38^, with peak values of *C*_*L*_ ≈ 5 achievable during a rapid pitch-up manoeuvre^28^. Nevertheless, stall cannot be delayed indefinitely, and will compromise control authority when it occurs^10,13,20,24^. This suggests a different performance objective, which is to minimize the distance flown under hazardous post-stall conditions.

We tested this hypothesis by identifying the transition point minimizing the distance flown at *C*_*L*_ > 4. This threshold was set high to avoid penalising the high lift coefficients that can be achieved transiently during a rapid pitch-up manoeuvre^28^, but the solution to the optimization proved robust to the selection of lower threshold lift coefficients (Ext. Fig. 3). The solution that minimizes stall distance is a deep swooping trajectory closely resembling that of an experienced bird (Fig. 3C). Neither the relative longitudinal position nor relative vertical position of the observed transition points differed significantly from the predicted optima in a linear mixed effects model fitting the combination of bird and perch spacing as a random effect (mean longitudinal deviation: -0.70%; 95% c.i.: -2.2%, 0.88%; mean vertical deviation: 0.47%; 95% c.i.: -0.028%, 0.97%), and the variation in both axes was comparable between (longitudinal s.d.: 3.0%; vertical s.d.: 1.0%) and within (longitudinal s.d.: 2.3%; vertical s.d.: 1.0%) groups. The model’s close quantitative fit to the data over a range of different perch spacings (Fig. 4) therefore supports our hypothesis that the hawks learned trajectories minimizing the distance flown under hazardous post-stall conditions.

**Figure 4.**
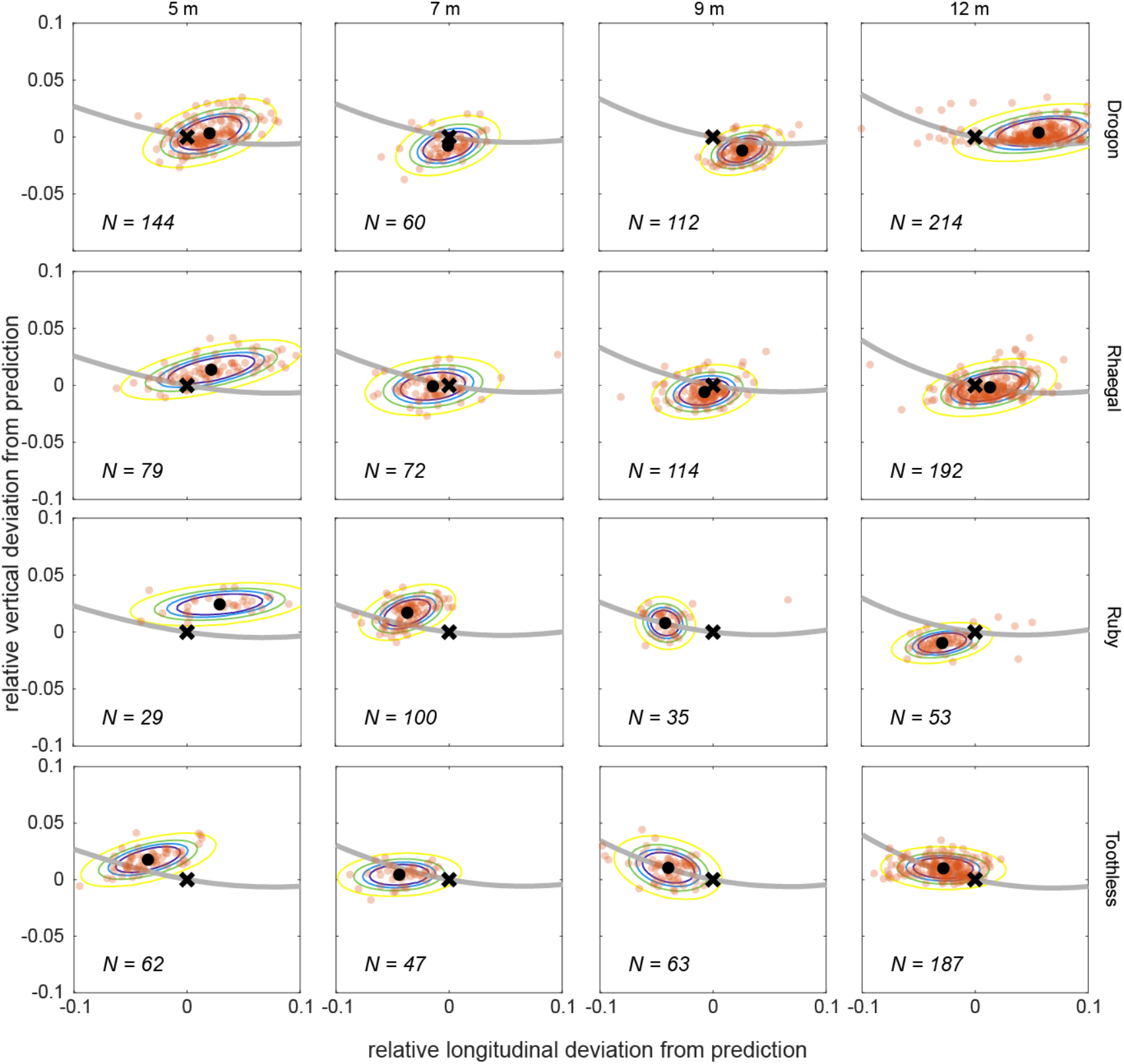
Fit of observed transition points to optima minimizing stall distance. At each combination of bird and perch spacing, the *N* observed transition points (opaque dots) are compared to the optimal transition point predicted to minimize stall distance under the model (black cross). The black dot denotes the sample mean for each test condition; coloured contours denote the 50^th^ to 95^th^ percentiles of a bivariate normal distribution fitted using the sample mean and covariance matrix within each group, at 5% intervals; grey line denotes the line of feasible transition points predicted under the model. Distance deviation is shown as a proportion of perch spacing distance.

How might this be implemented in practice? Controlling our two-phase model of perching amounts to launching at the optimal dive angle and flying at the optimal lift settings for a given perch spacing and chosen power output. It is an open question whether the birds learned to generalize from the specific perch spacing distances with which they were presented, but learning these parameter settings so as to minimize stall distance would presumably have entailed combining aeromechanical information from muscle^39^ or feather^1,2^ proprioceptors with distance information from static visual cues^5^ or optic flow^7,12,40^. Controlling these optimized parameter settings in open loop would be susceptible to external disturbances and internal error, but muscle power output is expected to be regulated locally under strain rate feedback from the muscle spindle cells^39^, and the lift force communicated to the body^41^ may be regulated locally under force feedback from the muscle Golgi tendon organs^39^. Analogous “fly-by-feel” concepts^42^ have demonstrated some notable successes in gust rejection in small air vehicles^43,44^, and may therefore prove important to controlling perching given the low-latency feedback that they provide^45,46^. Furthermore, given the physical significance of the transition point, which was always located at approximately 60% of perch spacing distance, it seems plausible that this learned optimum could have served as a virtual target for closed-loop trajectory control under visual feedback. This would be similar to the “entry gate” approach taken in one recent implementation of autonomous perching^18^, and could be important for correcting external perturbations before committing to the rapid pitch-up manoeuvre. In summary, a combination of visual and proprioceptive feedback is likely to be key to the learning and control of perching in birds, which points to the importance of combining both modalities in small air vehicles.

Above all, our modelling demonstrates that the usual currencies of time and energy that are typically optimized in steady flight behaviours^16^ are not necessarily minimized in the unsteady manoeuvres of large birds, which contrasts with previous findings from smaller birds in which energy is minimized^7^. Instead, our hawks learned swooping trajectories that minimized the distance flown under hazardous post-stall conditions, presumably reflecting the elevated hazard of landing at large size^6,11^. These findings have clear implications for deep reinforcement learning in autonomous systems, where identifying an appropriate cost function is critical^18^. Although stall may be important for generating high drag forces just before contact with the perch^10,13,18,20-24^, maintaining control authority is essential during any unsteady manoeuvre. We therefore propose that stall distance may prove a useful cost function for machine learning approaches aimed at acquiring robust perching capability on-the-fly, and envisage a new generation of bio-inspired autonomous vehicles that combine vision and mechanosensory feedback with deep reinforcement learning to achieve bird-like action intelligence.

## METHODS

### Experimental setup

We flew *n* = 4 captive-bred Harris’ Hawks (*Parabuteo unicinctus*) between two 1.25 m high A-frame perches positioned 5, 7, 9, or 12 m apart in a purpose-built motion capture studio (Fig. 1; Supplementary Movie 1). The sample comprised an experienced adult female (age: 7 years), and three inexperienced juvenile males (age: ≤ 0.5 years); see Table 1. The inexperienced birds had only previously flown during a brief period of fitness training conducted immediately before the experimental trials. The flights were undertaken in a windowless hall measuring 20.2 m x 6.1m, with a minimum ceiling height of 3.8 m and walls hung with camouflage netting to provide visual contrast. Flicker-free LED lights provided a mixture of direct 4000 K lighting and indirect 5000 K lighting at approximately 1000 lux, designed to mimic overcast morning or evening light.

### Experimental design

We collected trajectory data over an experimental period comprising 5-6 weeks of flight testing per bird. Each bird was flown individually between the perches, on a variable number of flights up to approximately 50 per day. The birds were motivated to fly from the take-off perch by the presentation of a small food reward on the landing perch, and usually responded immediately to this stimulus. The session was ended if the bird appeared tired or lacking in motivation, and the birds received a larger food reward at the end of the session. The birds were initially flown at 8 m perch spacing to introduce them to the testing environment and establish the measurement protocol. Perch spacing was then held at 12 m for approximately 2-3 weeks to allow us to identify when the birds behaviour had stabilized, before being randomized at 5, 7, or 9 m perch spacing daily thereafter. We treated the first 100 flights at 12 m perch spacing for each bird as comprising an initial familiarization period, by the end of which their flight behaviour had stabilized. The flights from this familiarisation period are not included in the main analysis, but are illustrated in Fig. 2A for the bird “Toothless”.

### Ethics statement

This work was approved by the Animal Welfare and Ethical Review Board of the Department of Zoology, University of Oxford, in accordance with University policy on the use of protected animals for scientific research, permit no. APA/1/5/ZOO/NASPA, and is considered not to pose any significant risk of causing pain, suffering, damage or lasting harm to the animals.

### Motion capture

We reconstructed the birds’ flight trajectories using a 20-camera motion capture system (Vicon Vantage 16, Vicon Motion Systems Ltd, Oxford, UK), mounted 3 m above the floor on scaffolding fixed around the walls. The motion capture system was turned on at least one hour before the start of the experiments, and was calibrated shortly before the first session (Vicon Active Calibration Wand), using Vicon Nexus 2 software for data acquisition. The motion capture cameras were set to record at 120 or 200 Hz under stroboscopic 850 nm infrared illumination, well outside the visible spectrum of these birds^47^, and a set of four high-definition video cameras (Vicon Vue) recorded synchronized reference video at 120 or 100 Hz, respectively. Each hawk was fitted with a rigid marker template comprising four 6.4 mm diameter spherical retroreflective markers (Fig. 1) worn on a falconry backpack secured by a pair of Teflon ribbons (TrackPack Mounting System, Marshall Radio Telemetry, UT, USA). The birds sometimes wore other retroreflective markers carried on fittings on the head, wings, or tail (see Supplementary Movie 1), but these are not included in the present analysis. A pair of 9.5 mm diameter spherical retroreflective markers was fixed to either end of each perch to identify the perch axis.

### Marker reconstruction

We used Vicon Nexus 2.7.6 software to reconstruct the positions of the markers within the flight volume, using a coordinate system corresponding to the principal axes of the flight hall. We removed any flights for which there were long sections of missing data, or for which the bird did not land at the perch, resulting in a sample of *n* = 1,585 complete flight trajectories suitable for analysis. This comprised *n* = 649 flights recorded at 12 m perch spacing, *n* = 324 flights at 9 m spacing, *n* = 279 flights at 7 m spacing, and *n* = 333 flights at 5 m spacing. The backpack and tail mount markers were usually visible on >70% of the recorded frames, but because of a challenging combination of dense marker placement, intermittent marker occlusion, and high-speed motion, the proprietary marker tracking algorithms were not uniformly successful in matching markers between frames. In addition, patches of specular reflection sometimes appeared as ghost markers. Consequently, although the Nexus software reconstructed the positions of all visible markers to a high degree of accuracy, it was not always able to label each marker reliably, or to identify every marker on every frame. We therefore wrote a custom script in MATLAB v2018a (The Mathworks Inc., MA, USA) which analysed the pattern of pairwise distances between markers in the rigid templates to label the anonymous markers.

### Marker labelling

The anonymous markers were labelled separately for each frame by using Procrustes analysis to match any visible markers to the known backpack template. We used the centroid of the resulting set of candidate backpack markers as an initial estimate of backpack position and fitted a quintic spline to interpolate its position on frames with missing data. We then used our initial or interpolated estimate of the backpack’s position on each frame to define a search volume matched to the size of the backpack template, and labelled any other markers falling within this search volume as candidate backpack markers. This two-stage labelling approach was able to accommodate missing markers and occasional ghost markers, and successfully identified the correct number of markers in >80% of all frames in which the backpack markers were visible. As the backpack sat directly between the scapulars, we took the centroid of the candidate backpack markers to approximate the position of the bird’s centre of mass, and estimated its velocity and acceleration by fitting and differentiating the smoothest quintic spline function passing through the positions measured on each frame.

### Trajectory analysis

Because the take-off and landing perches were located at the same height, every flight necessarily involved a powered flapping phase to overcome its drag losses. The three juvenile males flew directly between the perches by flapping on their first few flights (Fig. 2A), but quickly adopted the deep swooping trajectory typical of experienced birds including the adult female (Fig. 2B-E). This behaviour characteristically involves a powered dive followed by an unpowered climb (Fig. 1). Because the birds morphed smoothly between flapping and gliding flight, it was not possible to identify a unique point at which this transition occurred with reference to the wing kinematics. We instead identified the transition as occurring at the lowest point in the bird’s flight trajectory, which we determined using a band-stop Butterworth filter (2 to 19 Hz with maximum suppression at 6 Hz) to suppress body heave due to flapping.

### Take-off and landing

Each flight was initiated by a take-off jump during which the feet remained in contact with the perch. This jump ended with a jerk as the feet released from the perch, but the noise associated with the measured acceleration, particularly during flapping, makes it an unreliable marker of the onset of the flight phase. We therefore defined flight proper as beginning when the backpack reached a horizontal distance of 0.65 m from the take-off perch axis, this threshold distance being determined through visual inspection of the angular acceleration traces over many flights. The point of contact with the landing perch was likewise associated with a pronounced linear and angular acceleration, but for similar reasons we define the flight proper as ending when the backpack reached a horizontal distance of 0.35 m from the landing perch axis. The difference in these two threshold distances relates to the fact that the bird’s legs are extended caudad at take-off and ventrad upon landing. We found that the backpack was located approximately 0.30 m above the perch upon take-off, and approximately 0.15 m above the perch upon landing, which we used to define the initial and terminal conditions for the flight dynamics modelling.

### Flight dynamics model

To identify what flight performance objective(s) were optimized by the birds’ swooping trajectories, we built a simplified flight dynamics model for both flight phases (see Supporting Code). As a first-order modelling approach, we assumed constant aerodynamic lift (*L*) and power (*P*) on each phase, using settings *L* = *L*_dive_ and *P* = *P*_dive_ for the powered dive phase, and *L* = *L*_glide_ and *P* = 0 for the unpowered glide phase, where lift *L* is defined as the component of the aerodynamic force perpendicular to the flight path. Aerodynamic thrust (*T*) acting tangential to the flight path is modelled as *T* = *P/V* during the powered phase, where *V* is the bird’s airspeed neglecting any induced velocity component. This is opposed by aerodynamic drag (*D*), which we model as the sum of a lift-induced drag component and a combined parasite and profile drag component^32^:

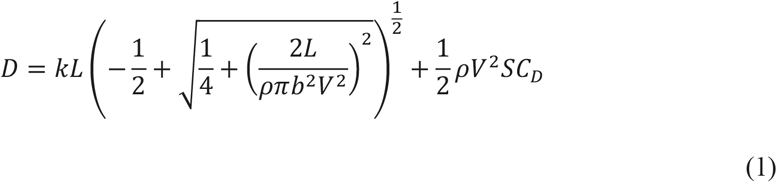

where 1 = 1.23 kg m^−3^ is air density, and where *b* is wingspan and *S* is wing area – both of which we assumed were maximal throughout the manoeuvre (Table 1). Eq. 1 models the induced drag in a generalized form that reflects how lift production is powered by the kinetic energy of the flow past the wings^48^. This expression results in high drag at low airspeeds, so although the angle of attack of the wing is not modelled explicitly in Eq. 1, the model implicitly captures the very high drag associated with operation at high angles of attack. We estimated the dimensionless induced drag factor *k* = 1.623 and combined profile and parasite drag coefficient *C*_*D*_ = 0.00994 by regressing the measured drag from an empirical glide polar for a Harris’ hawk^32^ against the predictors on the righthand side of Eq. 1.

With these assumptions, the rate of change in airspeed and flight path elevation angle (*γ*) can be expressed as:

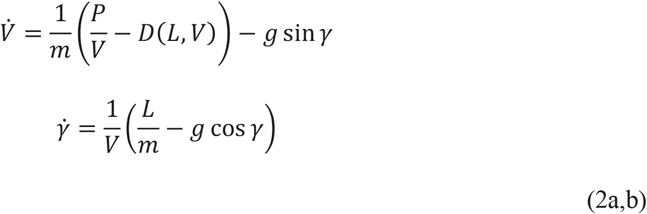

where *g* is gravitational acceleration and *m* is the bird’s mass. We modelled the resulting flight trajectories in lab-fixed Cartesian coordinates (*x,y*) by coupling Eqs. 1-2 for 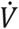 and 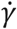 with the component kinematics equations:

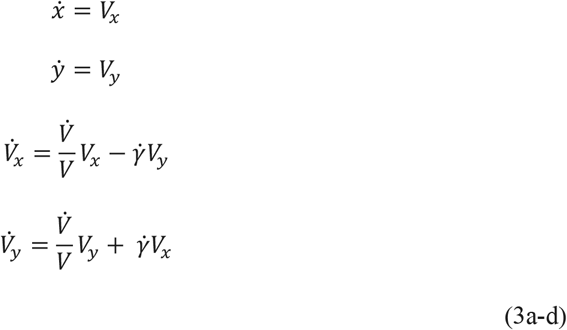

with 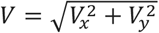. We integrated these ordinary differential equations numerically using the ode45 solver in Matlab, which is based on an explicit Runge-Kutta (4,5) formula, the Dormand-Price pair.

### Trajectory simulations

We simulated each bird individually to account for variation in flight morphology. We matched the initial speed *V*(0) of the simulations to the mean take-off speed 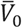 observed for each bird at the threshold horizontal distance of 0.65 m from the take-off perch (Table 1). We treated the initial dive angle *γ*(0) = *γ*_0_ as a free parameter, so the initial conditions for integrating Eqs. 3a-d were 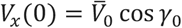 and 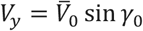 with *x*(0) = 0.65 m,, (0) = 1.55 m, and *γ*_0_ < 0 in the main text. We modelled the powered dive phase by assuming a fixed constant power setting *P* = *P*_dive_ in Eq. 2a, and by treating the constant lift setting in Eqs. 1-2a,b as a free parameter *L* = *L*_dive_. We optimized *P*_dive_ separately for each bird (see Main Text). With these assumptions, and for a given constant power setting *P*_dive_, each pair of free parameter settings {*γ*_0_, *L*_dive_} defines a unique flight trajectory for the powered flight phase. A subset of these powered dive trajectories passes through the horizontal, in the sense of having a point (*x*_T_, *y*_T_) at which *V*. = 0 with *V*_*T*_ = *V*_*x*_ ≠ 0. This subset defines the set of reachable combinations of position and speed at which the transition from powered dive to unpowered glide can occur under the model.

The initial conditions for the glide phase are given by the position (*x*_T_, *y*_T_) and velocity *V*(_T_, 0) of the bird at this transition point. Each pair of parameter settings {*γ*_0_, *L*_dive_} for the powered dive phase therefore defines a family of possible flight trajectories for the glide phase, parameterized only by its constant lift setting *L* = *L*_glide_. Hence, for any given pair of parameter settings {*γ*_0_, *L*_dive_}, we are left only to solve for the unique value of *L*_glide_ that will produce a trajectory intercepting the point of contact with the landing perch at *x* = *s* − 0.35 m and *y* = 1.40 m, where *s* is the perch spacing (see above). In practice, there are only certain combinations of {*γ*_0_, *L*_dive_} that will bring the simulated bird to the landing perch at a realistic speed, so it proved most efficient to solve the glide phase backwards in time from the point of contact with the landing perch, and to match the solutions for the two flight phases at the transition point (see Supporting Code). We therefore fixed the initial speed of this backwards simulation of the glide phase to the mean landing speed 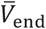 observed for each at the threshold horizontal distance of 0.35 m from the landing perch (Table 1). For the purposes of finding matching solutions, we treated both the flight path angle at the point of contact (*γ*_end_) and the constant lift setting for the glide phase (*L*_glide_) as free parameters. These parameters {*γ*_end_, *L*_glide_} then become fixed for a given pair of parameter settings {*γ*_0_, *L*_dive_} once the matching solution for the powered dive phase is found.

### Feasible trajectory search

We identify feasible trajectories as those which bring an individual bird to its landing perch at the same mean speed and position as we observed in the experiments. For a given constant power setting *P*_dive_, this constraint defines a line of feasible transition points corresponding to a line of feasible parameter settings {*γ*_0_, *L*_dive_}. We implemented the search for feasible transition points as a constrained minimization problem solved using an interior-point algorithm in MATLAB 2020a. We constrained the difference in transition point position (*x*_T_, *y*_T_) and velocity *V*(_T_, 0) between the end of the powered phase and start of the unpowered phase to be zero, and first solved for the parameter settings {*γ*_0_, *L*_dive_} and {*γ*_end_, *L*_glide_} that would have placed the transition point at the landing perch. We then took these parameter settings as initial values when solving for the parameter settings that would have caused the transition point to be placed a small increment ahead of the perch, which we set as the target of the minimization. We repeated this process to place the transition point another small increment in distance ahead of the perch, inheriting the parameter settings of the previous solution as initial values for the next round, until the complete line of feasible transition points had been found (see Ext. Fig. 4 for a detailed summary of solutions). It is important to note that other transition points falling close to this line could also be physically feasible in the sense of bringing the bird to the landing perch, but these will be associated with higher or lower speeds than those observed at the point of contact.

### Trajectory optimization

The unique mapping that exists between the parameter settings {*γ*_0_, *L*_dive_} and transition point {*x*_T_, *y*_T_, *V*_T_} means that any property of a given flight trajectory is also a property of its transition point. This includes the duration (*τ*) and energetic cost (*E*) of the flight, and the distance from the landing perch at which stall occurs (*d*_stall_) – each of which may be considered a candidate objective function for minimization. We identified the optimal transition point at which each of these objectives was minimized by a direct search along the line of feasible transition points. Under the two-phase model of perching, the duration of a flight trajectory is implicit in its solution as *τ* = *τ*_dive_ + *τ*_glide_. Minimizing the total flight duration *τ* therefore entails jointly minimizing the duration of the powered dive phase *τ*_dive_ and unpowered glide phase *τ*_glide_. In contrast, given the constant power assumption, the energetic cost of a flight trajectory is simply *E* = *P*_dive_*τ*_dive_, so for a given constant power setting *P*_dive_, minimizing the energetic cost of the flight is equivalent to minimizing the duration of the powered dive phase *τ*_dive_ alone. Wing stall is a complex phenomenon, so we did not model its effects directly. However, because lift varies as *L* = *ρV*^2^*SC*_*L*_*/*2, stall is implicit in the very high values of the lift coefficient *C*_*L*_ that are needed to meet the constant lift requirement *L* = *L*_glide_ at the low speeds *V* reached as the bird decelerates on approach to the perch. Minimizing the distance flown post-stall therefore amounts to penalising flight at values of *C*_*L*_ exceeding some specified threshold, which we implemented by minimizing the distance flown at *C*_*L*_ > 4 (see Main Text). The predicted location of the optimal transition point was robust to this choice of threshold, moving ≤ 1.5*γ* of perch spacing distance per unit decrement in the threshold value of *C*_*L*_ (Ext. Fig. 3).

### Statistics

We modelled the longitudinal and vertical position of the observed transition points as a proportion of perch spacing distance using a linear mixed effects model (`fitlme`) in MATLAB 2020a. We treated the mean centred perch distance as a covariate, and individual as a random effect, such that: `RelativePosition ∼ 1 + MeanCentredPerchDistance + (1+MeanCentredPerchDistance|BirdID`), using two-tailed *p*-values for statistical inference. The model for relative longitudinal position identified 22 of the 1,585 observed transition points as outliers, with residuals >3 times the residual standard deviation. Two of these outliers were attributable to motion capture error; the remainder corresponded to non-swooping flight behaviours (7 flights) and/or trajectories at 5 m perch spacing for which the transition occurred at a local rather than global minimum in height (17 flights, of which 11 were associated with the adult bird ‘Ruby’). As these outliers are unrepresentative of the swooping behaviour analysed in this paper, we excluded them from this and subsequent analyses, leaving a slightly smaller sample of 1,563 flights (see Supporting Code).

To test how well the flight dynamics model predicted the observed transition points, we fitted the longitudinal and vertical distances between the predicted optima and the observed transition points using a linear mixed effects model. We treated the combination of perch distance and individual as a random effect to account for the single prediction for each experimental condition, such that: `Distance ∼ 1 + (1|PerchDistance:BirdID)`. In this model, a significant intercept indicates a systematic deviation between the observed and predicted transition points. We computed the sample mean and covariance matrix of the longitudinal and vertical position of the observed transition points at every combination of individual identity and perch distance, and generated the percentiles of the corresponding bivariate normal distribution by computing their squared Mahalanobis distance from the mean using the Chi-square distribution on 2 degrees of freedom.

## Acknowledgments

This project has received funding from the European Research Council (ERC) under the European Union’s Horizon 2020 research and innovation programme (Grant Agreement No. 682501). LF’s work was supported by funding from the Biotechnology and Biological Sciences Research Council (BBSRC) [grant number BB/M011224/1], via the Interdisciplinary Bioscience Doctoral Training Partnership. The authors thank Emma Borsier, Natalia Pérez-Campanero, and Sofía Miñano-González for comments, and Mark Parker and Lucy Larkman for animal husbandry and handling during experiments.

## Author contributions

**MKH, LAF** Conceptualization, Methodology, Software, Investigation, Formal analysis, Data curation, Writing – original draft, Writing – review & editing, Visualization. **CHB** Methodology, Investigation, Writing – review & editing. **GKT** Conceptualization, Methodology, Investigation, Formal analysis, Resources, Writing – original draft, Writing – review & editing, Supervision, Project administration, Funding acquisition.

## Competing interests

The authors declare no competing interests.

## Additional information

Supplementary Information is available for this paper. Correspondence and requests for materials should be addressed to.

## Data and code availability

The motion capture data and code that support the findings of this study are available in figshare with the identifier doi:10.6084/m9.figshare.16529328

**Extended Data Figure 1.**
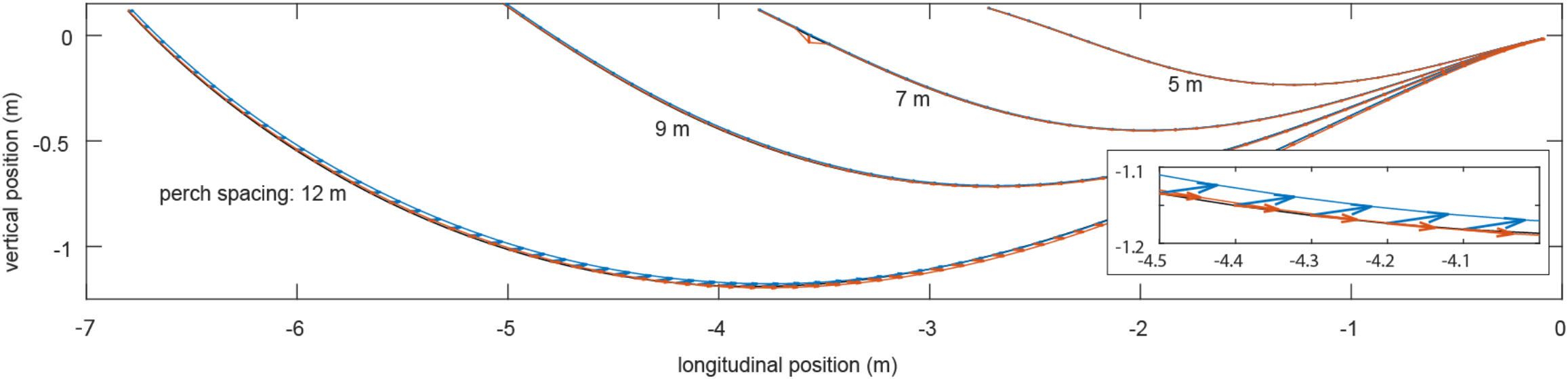
Impact of ground effect on the line of feasible transition points. We assessed the impact of ground effect by re-evaluating the powered dive phase (red) and unpowered glide phase (blue) with an aerodynamic model of ground effect^34^ included for the same set of feasible parameter settings as we had identified previously without ground effect (black). Whereas the powered dive transitions smoothly into the unpowered glide for the original solutions without ground effect (black), the introduction of ground effect without any changes to the model parameters necessarily results in a small mismatch between the two flight phases. The extent of this mismatch is shown by the arrows plotting the change in position of the transition point predicted at the end of the powered dive (red arrows) and the start of the unpowered glide (blue arrows) for trajectories satisfying the take-off and landing constraints. The effect, shown here for the juvenile bird “Drogon”, is marginal; the enlarged inset shows the effect on transition points close to the ground at 12 m perch spacing for which the impact of ground effect is greatest.

**Extended Data Figure 2.**
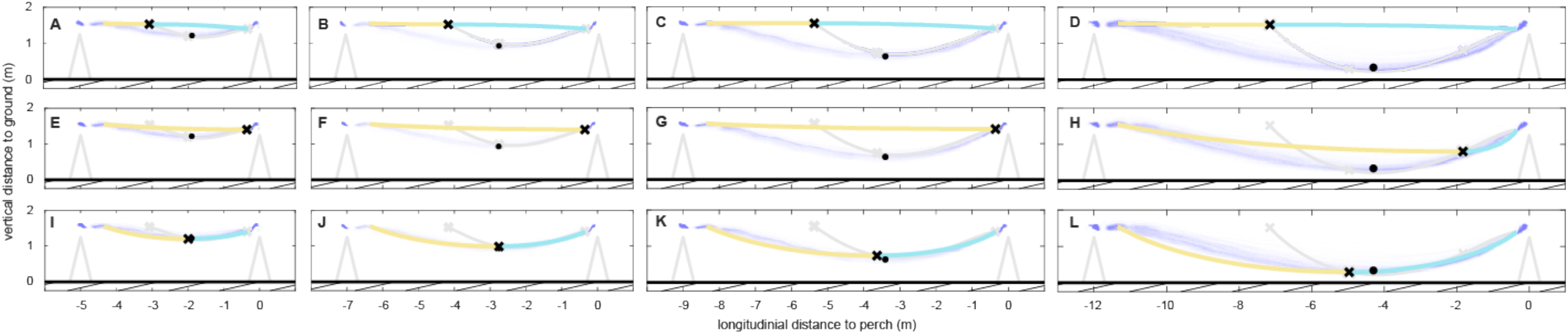
Optimal perching trajectories minimizing different cost functions at different perch spacings. Thick coloured lines show trajectories predicted for the juvenile bird “Drogon” at 5 m, 7 m, 9 m, or 12 m perch spacing under the two-phase perching model, comprising a powered dive (yellow line), transitioning into an unpowered glide (cyan line), and minimizing: **(A-D)** energetic cost; **(E-H)** flight duration; or **(I-L)** stall distance. The location of the optimal transition point (black cross) along the line of feasible transition points (grey line) is only close to the mean observed transition point (black dot) if stall distance is optimized (**I-L**); observed trajectories are shown as a spatial histogram (lilac shading).

**Extended Data Figure 3.**
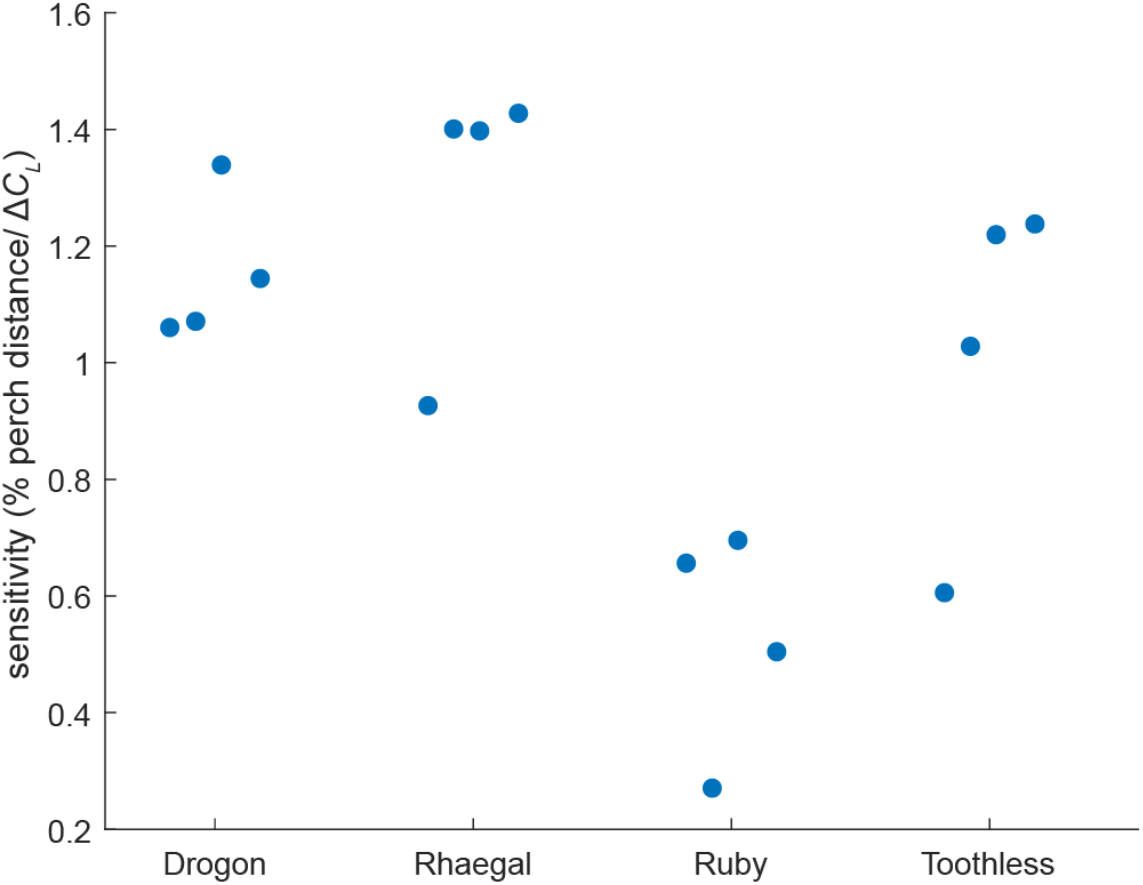
Effect of penalty threshold on location of stall-optimal transition points. In the main text, the location of the optimal transition point minimizing stall distance is determined by minimizing the distance flown at lift coefficients *C*_*L*_> 4. Decreasing this penalty threshold causes the optimum to move closer to the landing perch along the line of feasible transition points, eventually reaching a point of minimum distance from the landing perch at some specific decrement Δ*C*_*L*_, depending on individual bird and perch spacing. We quantified the sensitivity of the solution to the choice of penalty threshold by expressing the maximal displacement of the optimal transition point along the line of feasible transitions points relative to the corresponding decrement in penalty threshold Δ*C*_*L*_. The points for each bird represent 5 m, 7 m, 9 m, and 12 m perch spacing, and the displacement of the optimal transition point is normalized by perch spacing distance. Note the robustness of the location of the stall-optimal transition point to the choice of penalty threshold (displacement sensitivity <1.5% of perch spacing distance per unit decrement in the penalty threshold value of *C*_*L*_).

**Extended Data Figure 4.**
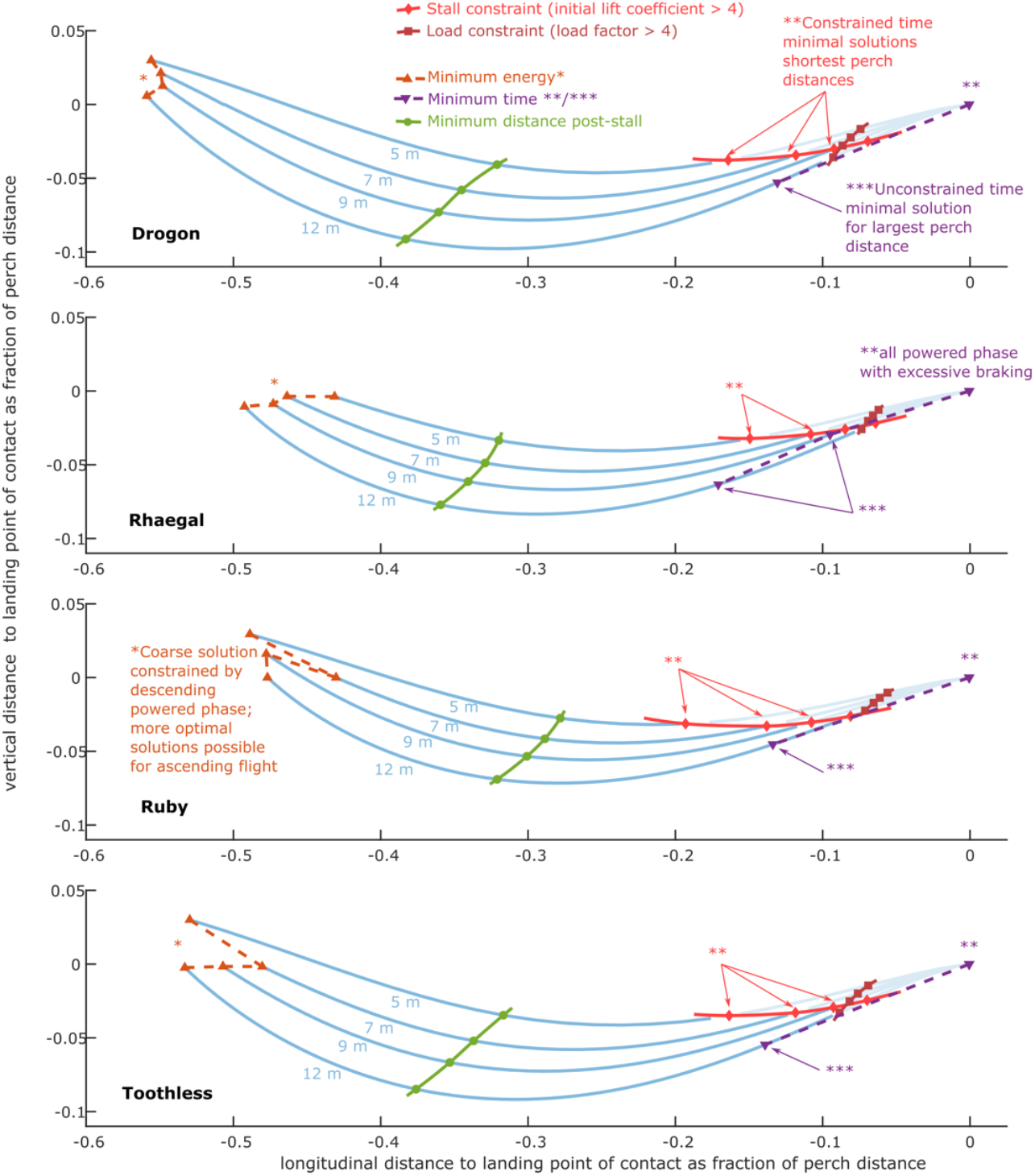
Effect of constraints on location of feasible transition points. Blue lines plot the set of feasible transition points for each bird and perch spacing, normalized relative to perch spacing distance. Green dots plot stall-optimal solutions minimizing the distance flown at *C*_*L*_> 4. Dark red triangles show energy-minimizing solutions, allowing the possibility of an initial climb at *γ*_0_ > 0, which can sometimes be more efficient than diving at *γ*_0_ ≤ 0 as observed and assumed in the main text (see Methods). Purple triangles plot time-optimal solutions, which can involve very high load factors (*L/*(*mg*)) ≥ 4) and very high lift coefficients (*C*_*L*_≥ 4) from the transition point onward. Dark red squares and bright red diamonds show constrained time-optimal solutions subject to the inequality constraints *L/*(*mg*) < 4 or *C*_*L*_< 4, respectively, at the start of the glide phase.

**Supplementary Movie 1. Reference video of typical swooping perching trajectory**. Reference video showing a sequence of flights at 12 m perch spacing by juvenile bird “Drogon”. In this sequence, the bird is seen wearing a falconry backpack, a falconry tail mount, and a falconry hood that had been modified to avoid obscuring the bird’s vision. Retroreflective markers of various sizes are visible attached to the backpack, hood, tail mount, and wing feathers, but only the 6.4 mm diameter backpack markers were tracked for the purposes of this study. The video images have been undistorted automatically using the camera calibration from the motion capture system, and the falconer’s face has been anonymised, but no other image manipulations have been applied. Backpack marker paths tracked retrospectively by the Vicon Nexus software are overlain ahead of the trajectories that the markers follow.

## Notes

https://doi.org/10.6084/m9.figshare.16529328

### Competing Interest Statement

The authors have declared no competing interest.

### Summary of Updates

Added co-author and endnotes; implemented improved method of trajectory smoothing resulting in some small changes to outlier exclusion and sample size; made extensive small improvements throughout text and figures; added supplemental files.

## References

1 Carruthers, A. C., Thomas, A. L. R. & Taylor, G. K. Automatic aeroelastic devices in the wings of a steppe eagle Aquila nipalensis. J Exp Biol 210, 4136–4149, doi:10.1242/jeb.011197 (2007).

2 Carruthers, A. C., Thomas, A. L. R., Walker, S. M. & Taylor, G. K. Mechanics and aerodynamics of perching manoeuvres in a large bird of prey. Aeronautical Journal 114, 673–680, doi:10.1017/s0001924000004152 (2010).

3 Green, P. R. & Cheng, P. Variation in kinematics and dynamics of the landing flights of pigeons on a novel perch. Journal of Experimental Biology 201, 3309–3316 (1998).

4 Berg, A. M. & Biewener, A. A. Wing and body kinematics of takeoff and landing flight in the pigeon (Columba livia). J Exp Biol 213, 1651–1658, doi:10.1242/jeb.038109 (2010).

5 Kress, D., van Bokhorst, E. & Lentink, D. How Lovebirds Maneuver Rapidly Using Super-Fast Head Saccades and Image Feature Stabilization. PLoS One 10, e0129287, doi:10.1371/journal.pone.0129287 (2015).

6 Polet, D. T. & Rival, D. E. Rapid area change in pitch-up manoeuvres of small perching birds. Bioinspir Biomim 10, 066004, doi:10.1088/1748-3190/10/6/066004 (2015).

7 Chin, D. D. & Lentink, D. How birds direct impulse to minimize the energetic cost of foraging flight. Science Advances 3, doi:10.1126/sciadv.1603041 (2017).

8 Quinn, D. et al. How lovebirds maneuver through lateral gusts with minimal visual information. Proc Natl Acad Sci U S A 116, 15033–15041, doi:10.1073/pnas.1903422116 (2019).

9 Pennycuick, C. J. Mechanics of bird migration. Ibis 111, 525-+, doi:10.1111/j.1474-919X.1969.tb02566.x (1969).

10 Wickenheiser, A. M. & Garcia, E. Optimization of Perching Maneuvers Through Vehicle Morphing. Journal of Guidance, Control, and Dynamics 31, 815–823, doi:10.2514/1.33819 (2008).

11 Maynard Smith, J. The importance of the nervous system in the evolution of animal flight. Evolution 6, 127–129, doi:10.2307/2405510 (1952).

12 Lee, D. N., Davies, M. N. O., Green, P. R. & Van der Weel, F. R. R. Visual control of velocity of approach by pigeons when landing. Journal of Experimental Biology 180, 85–104 (1993).

13 Moore, J., Cory, R. & Tedrake, R. Robust post-stall perching with a simple fixed-wing glider using LQR-Trees. Bioinspir Biomim 9, 025013, doi:10.1088/1748-3182/9/2/025013 (2014).

14 Zhang, Z., Xie, P. & Ma, O. Bio-Inspired Trajectory Generation for UAV Perching Movement Based on Tau Theory. International Journal of Advanced Robotic Systems 11, doi:10.5772/58898 (2014).

15 Chirarattananon, P., Ma, K. Y. & Wood, R. J. Perching with a robotic insect using adaptive tracking control and iterative learning control. The International Journal of Robotics Research 35, 1185–1206, doi:10.1177/0278364916632896 (2016).

16 Hedenstrom, A. & Alerstam, T. Optimal flight speed of birds. Philosophical Transactions of the Royal Society of London Series B-Biological Sciences 348, 471–487, doi:10.1098/rstb.1995.0082 (1995).

17 Mills, R., Hildenbrandt, H., Taylor, G. K. & Hemelrijk, C. K. Physics-based simulations of aerial attacks by peregrine falcons reveal that stooping at high speed maximizes catch success against agile prey. PLoS Comput Biol 14, e1006044, doi:10.1371/journal.pcbi.1006044 (2018).

18 Waldock, A., Greatwood, C., Salama, F. & Richardson, T. Learning to Perform a Perched Landing on the Ground Using Deep Reinforcement Learning. Journal of Intelligent & Robotic Systems 92, 685–704, doi:10.1007/s10846-017-0696-1 (2017).

19 Novati, G., Mahadevan, L. & Koumoutsakos, P. Controlled gliding and perching through deep-reinforcement-learning. Physical Review Fluids 4, doi:10.1103/PhysRevFluids.4.093902 (2019).

20 Cory, R. & Tedrake, R. in AIAA Guidance, Navigation & Control Exhibit (American Institute of Aeronautics and Astronautics, Honolulu, Hawaii, 2008).

21 Lussier Desbiens, A., Asbeck, A. T. & Cutkosky, M. R. Landing, perching and taking off from vertical surfaces. The International Journal of Robotics Research 30, 355–370, doi:10.1177/0278364910393286 (2011).

22 Paranjape, A. A., Chung, S.-J. & Kim, J. Novel Dihedral-Based Control of Flapping-Wing Aircraft With Application to Perching. IEEE Transactions on Robotics 29, 1071–1084, doi:10.1109/tro.2013.2268947 (2013).

23 Greatwood, C., Waldock, A. & Richardson, T. Perched landing manoeuvres with a variable sweep wing UAV. Aerospace Science and Technology 71, 510–520, doi:10.1016/j.ast.2017.09.034 (2017).

24 Roderick, W. R., Cutkosky, M. R. & Lentink, D. Touchdown to take-off: at the interface of flight and surface locomotion. Interface Focus 7, 20160094, doi:10.1098/rsfs.2016.0094 (2017).

25 Provini, P., Tobalske, B. W., Crandell, K. E. & Abourachid, A. Transition from wing to leg forces during landing in birds. J Exp Biol 217, 2659–2666, doi:10.1242/jeb.104588 (2014).

26 Crandell, K. E., Smith, A. F., Crino, O. L. & Tobalske, B. W. Coping with compliance during take-off and landing in the diamond dove (Geopelia cuneata). PLoS One 13, e0199662, doi:10.1371/journal.pone.0199662 (2018).

27 Roderick, W. R., Chin, D. D., Cutkosky, M. R. & Lentink, D. Birds land reliably on complex surfaces by adapting their foot-surface interactions upon contact. Elife 8, doi:10.7554/eLife.46415 (2019).

28 Polet Delyle T., Rival David E. & Weymouth Gabriel D. Unsteady dynamics of rapid perching manoeuvres. Journal of Fluid Mechanics 767, 323–341, doi:10.1017/jfm.2015.61 (2015).

29 Tang, D. et al. Shape reconstructions and morphing kinematics of an eagle during perching manoeuvres. Chinese Physics B 29, doi:10.1088/1674-1056/ab610a (2020).

30 Wang, Y., Tobalske, B. W., Cheng, B. & Deng, X. Gravitation-enabled Forward Acceleration during Flap-bounding Flight in Birds. Journal of Bionic Engineering 15, 505–515, doi:10.1007/s42235-018-0041-9 (2018).

31 Goldstein, H., Poole, C. & Safko, J. Classical Mechanics. 3rd edn, (Addison Wesley, 2002).

32 Tucker, V. A. & Heine, C. Aerodynamics of gliding flight in a Harris’ Hawk, Parabuteo unicintus. Journal of Experimental Biology 149, 469–489 (1990).

33 Song, J. Fly low: The ground effect of a barn owl (Tyto alba) in gliding flight. Proceedings of the Institution of Mechanical Engineers, Part C: Journal of Mechanical Engineering Science 235, 308–318, doi:10.1177/0954406220943939 (2020).

34 Rayner, J. M. V. On the aerodynamics of animal flight in ground effect. Philosophical Transactions of the Royal Society of London. Series B: Biological Sciences 334, 119–128, doi:10.1098/rstb.1991.0101 (1991).

35 Pennycuick, C. J., Fuller, M. R. & McAllister, L. Climbing performance of Harris’ Hawks (Parabuteo unicinctus) with added load – implications for muscle mechanics and for radiotracking. Journal of Experimental Biology 142, 17–29 (1989).

36 Van Walsum, T. A. et al. Exploring the relationship between flapping behaviour and accelerometer signal during ascending flight, and a new approach to calibration. Ibis 162, 13–26, doi:10.1111/ibi.12710 (2019).

37 Reich, G. W., Eastep, F. E., Altman, A. & Albertani, R. Transient Poststall Aerodynamic Modeling for Extreme Maneuvers in Micro Air Vehicles. Journal of Aircraft 48, 403–411, doi:10.2514/1.C000278 (2011).

38 Uhlig, D. V. & Selig, M. S. Modeling Micro Air Vehicle Aerodynamics in Unsteady High Angle-of-Attack Flight. Journal of Aircraft 54, 1064–1075, doi:10.2514/1.C033755 (2017).

39 Granatosky, M. C. et al. Variation in limb loading magnitude and timing in tetrapods. J Exp Biol 223, doi:10.1242/jeb.201525 (2020).

40 Davies, M. N. O. & Green, P. R. Optic flow-field variables trigger landing in hawk but not in pigeons. Naturwissenschaften 77, 142–144, doi:10.1007/bf01134481 (1990).

41 Reynolds, K. V., Thomas, A. L. & Taylor, G. K. Wing tucks are a response to atmospheric turbulence in the soaring flight of the steppe eagle Aquila nipalensis. J R Soc Interface 11, 20140645, doi:10.1098/rsif.2014.0645 (2014).

42 Yeo, D., Atkins, E. M., Bernal, L. P. & Shyy, W. Fixed-Wing Unmanned Aircraft In-Flight Pitch and Yaw Control Moment Sensing. Journal of Aircraft 52, 403–420, doi:10.2514/1.C032682 (2015).

43 Araujo-Estrada, S. A. et al. in AIAA Guidance, Navigation, and Control Conference (2017).

44 Castano, L., Airoldi, S., McKenna, T. & Humbert, J. S. in 14th AIAA Aviation Technology, Integration, and Operations Conference (American Institute of Aeronautics and Astronautics, Atlanta, GA, 2014).

45 Thompson, R. A., Evers, J. H. & Stewart, K. C. in AIAA Atmospheric Flight Mechanics Conference (American Institute of Aeronautics and Astronautics, Toronto, Ontario, Canada, 2010).

46 Gremillion, G. M., Castano, L. & Humbert, J. S. in AIAA Guidance, Navigation, and Control Conference (American Institute of Aeronautics and Astronautics, Kissimmee, FL, 2015).

47 Potier, S., Mitkus, M. & Kelber, A. High resolution of colour vision, but low contrast sensitivity in a diurnal raptor. Proc. R. Soc. B 285, doi:10.1098/rspb.2018.1036 (2018).

48 Seddon, J. M. N. S., Basic helicopter aerodynamics. (Wiley, 2011).

